# The *osm-9* mutant strain is defective in the recovery phase of long-term memory dynamics

**DOI:** 10.64898/2025.12.30.697040

**Authors:** Nikki I. Ma, Fatema M. Saifuddin, Nando Munoz Lobato, Noelle D. L’Etoile

## Abstract

Memory by definition is the storage of learned information from experiences where it can be recalled in the future. Memory is essential for organisms of any level, from single celled bacteria to complex mammals like whales in order for them to adapt and survive. The storage of long term memory is known to start in the hippocampus but long-term memory is not simply contained in this one region in the brain but across multiple regions including the cortex, amygdala and nucleus accumbens. This diversity of locations that memory can be located in prompts the question of what molecular components within the nervous system are required to establish and maintain long term memory. To dive into this, we utilize the simple neuroanatomy of *C. elegans* with only 302 neurons to examine the effects of mutations of specific genes on the success of forming long term memories. Previous work established that *C. elegans* with defective OSM-9 proteins are unable to sustain an aversion to butanone (an odor that they are innately attracted to) after it is paired once with a negative stimulus. This helped established the importance of OSM-9 proteins for short term memory but how can the OSM-9 protein – a homolog to human TRPV5 and TRPV6 commonly associated with recognising pain and thermoregulation rather than memory – play a part within the dynamics of space training induced long term memory? Here we show that the loss of the OSM-9 protein affects the dynamics of memory consolidation, which is the process by which short term memory is converted to long term memory. Previously, OSM-9 protein was implicated in the formation of short term memory which was termed adaptation. Here, we examine spaced training induced memory that is resistant to re-feeding and depends on sleep. This study indicates that after spaced training, the OSM-9 protein is required for the conversion of short term memory to long term memory and not acquisition. Furthermore, we confirmed that 30 minutes on food after spaced training, Wild type *C. elegans* seem to lose their memory but gain it back 120 minutes after, emphasizing the dynamics of memory and revealing that it is not a smoothly continuous process. Our results demonstrate that TRPV containing channels plays a role in stabilizing memory for long term use. Analyzing the specific point in which the OSM-9 protein plays a part in reactivating memory may help pave the way to a more detailed understanding of memory dynamics and the processes involved in recovering memory. Future work includes: identifying where the loss of memory goes, whether its masked or truly disappears can further explain what the OSM-9 protein supports when long term memory is formed; understanding if OSM-9 is directly involved with the process of “reviving” the memory or is it involved in the development of the *C. elegans* circuit which allows for this process to occur.

## Introduction

Memory is required for animals to learn from their experiences; if animals can not remember how to change their behavior based on past mistakes, they are at risk of reducing their fitness. Thus, memory is under strong evolutionary selection and is exhibited by organisms ranging from single celled Stentor (Rajan and Marshall 2025 Current Biology) to drosophila larvae (Lesar et al., 2021) to humans (Kandell 2014). An understanding of the mechanisms by which organisms that are quite diverged evolutionarily learn and commit what they have learned to memory have the potential to either reveal new biology if it differs or uncover some of the very basis by which human memory is formed if it is conserved. Previously, we discovered that the 302-neuron *C. elegans* nervous system requires sleep to consolidate an aversive memory of being starved (Chandra, Farah and Munoz Lobato, 2023). This is the simplest nervous system known to produce sleep-dependent memory.

We made this finding when we examined memory after 30-minute intervals of recovery on food and noticed that memory seemed to disappear in the first 30 minutes and reappear by 120 minutes of feeding. Close examination of the animals revealed that they were asleep during that time during which memory was labile. The labile memory’s dependence on sleep mirrors human memory consolidation (Kandell 2014). Though we have a wealth of information about the genes required for memory formation and some of the brain regions engaged (Ortega de San Luis, Ryan, 2022), the complexity of the mammalian brain has obscured in depth examination of the exact cellular locus for each stage of memory. Puzzlingly, we know that initial memory requires the hippocampus but long lasting memories are stored throughout the cortex. How memory transitions from one locus to the next and how it becomes more enduring in the process is just not understood (Kendall, 2014.

We turned to the completely transparent *C. elegans*, with many mutants that affect memory and a very simple neuroanatomy consisting of a mere 302 neurons to examine how a short term memory is consolidated. Its simple, optically accessible anatomy, allows for easy interrogation of the cellular locus for short and long term memory. Furthermore, its robust genetics has allowed other labs to identify mutants that fail to either learn or remember. Previous screens found that worms defective in the *osm-9(ky10)* gene that codes for the TRPV (transient potential vanilloid channel) OSM-9 protein – a homolog to human TRPV5 and TRPV6 – are unable to adapt to the odor butanone, an odor that *C. elegans* are innately attracted to (Colbert and Bargmann, Neuron 1995). Prior work shows that *osm-9(ky10)* plays a role in short term memory (Benedetti et al., 2022 BioRxiv). However, OSM-9 is not required for sleep as our recent work shows that mutants lacking the OSM-9 protein sleep as well as wildtypes (Benedetti et al., 2022 BioRxiv)..

Benedetti et al., shows that OSM-9 is expressed in four sensory neurons and that loss of expression within these neurons leads to loss of long-term memory. These cells are: the AWA (responds to attractive food-associated odors), ADL (responds to pheromones and food), ADF (responds to food and infection) and ASH (responds to noxious stimuli) neurons.

Here, we show that the *osm-9(ky10)* gene is not required for learning or the initial apparent loss of memory in the first 30 minutes of recovery on food but that it is required for memory to reappear between 30 and 120 minutes on food. This study paves the way for a more detailed analysis of neural dynamics, cellular and circuit processes during the recovery phase that can now be focused on the four cells, ADL, ADF, ASH and AWA that are required to express OSM-9 in order for animals to consolidate long term olfactory memory.

## Results

To understand the role of the OSM-9 protein in the formation of long-term memory, we introduce an aversive memory through classical conditioning (Figure 1A). Specifically, four Larval stage 4 (L4) worms are placed onto a growth plate (Figure 1A,1) where they lay around 200 eggs which will be used in the experiment once they reach their first day of adulthood. Worms are washed from their growth plates, they are then separated into two training groups in 1.5 ml microfuge tubes that contain either buffer (control) or butanone (odor) diluted in buffer (Figure 1A, 2). The animals that are exposed to butanone without food lose their innate attraction to that odor while the control group that is exposed just to buffer while starving do not. After 80 minutes of starvation, they are placed within a food solution (which does not have butanone) for 30 minutes, this ensures that the *C. elegans* are still sustained and reinforces that butanone is not a food-associated odor. After, they are then rotated in their specified training solutions for a total of two cycles.

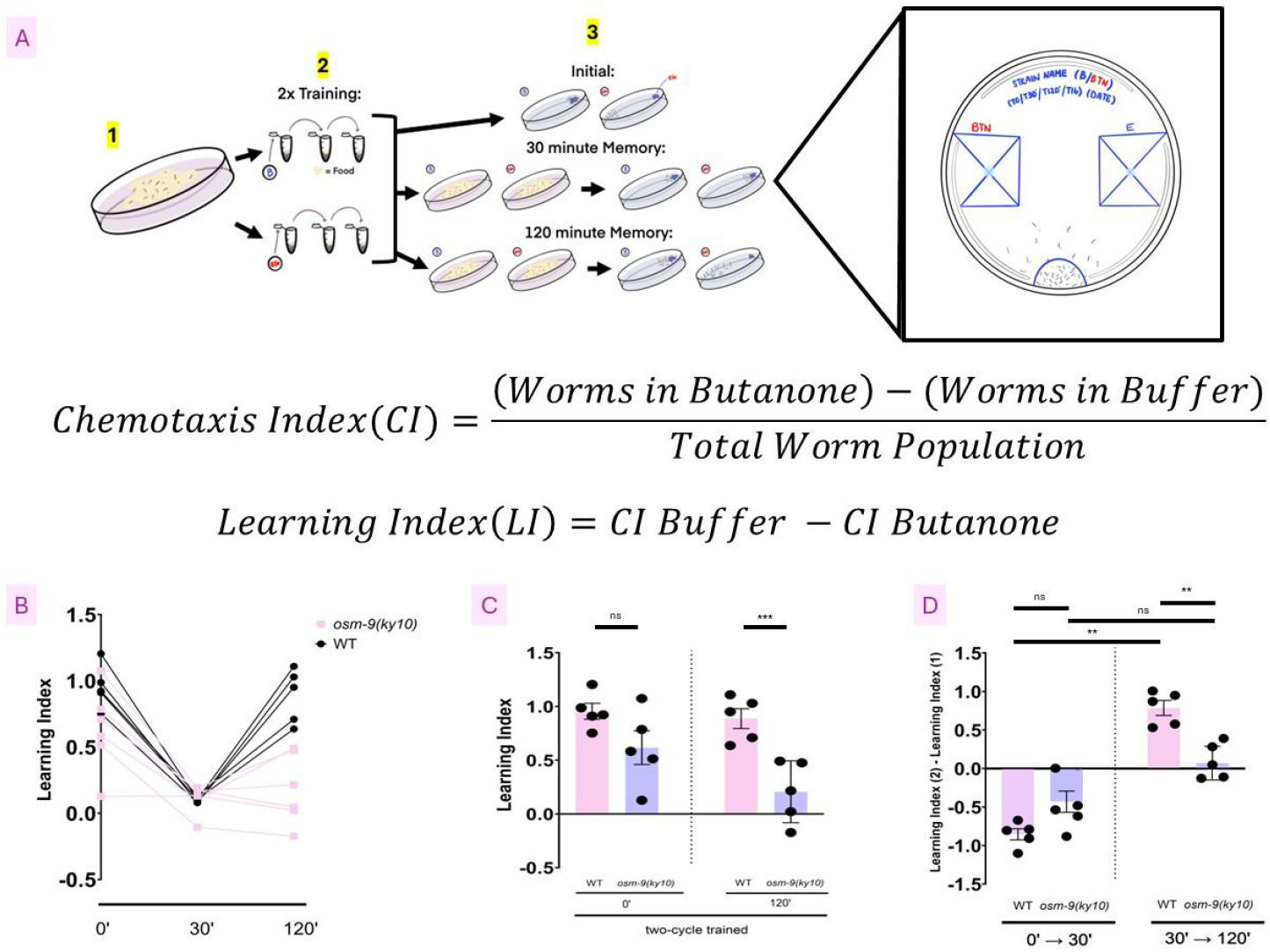
Figure 1a. Schematic of two cycle odor-conditioning and spaced training paradigm Unstarved populations of *C. elegans* are washed from their growth plates before being separated into two training groups in 1.5 ml tubes: buffer control (top) and butanone odor (bottom). Worms are rotated in liquid while being exposed to either buffer (control) or butanone (odor) for 80 minutes then washed and placed in liquid food for 30 minutes. The training with buffer or butanone was then repeated for a total of two training sessions. After the last training, each population is split into three. The first is subjected to a chemotaxis assay to butanone (top) at time 0’. The next is placed on plates containing food and left for 30 minutes before being assayed (middle), or a food plate for 120 minutes before being assayed (bottom). Cartoons of the chemotaxis plates are presented in the zoom box. The population is placed within the semi-circle at the bottom of the petri dish and allowed to run across the dish that has diluted butanone in the right box and the diluent ethanol in the left box (refer to the cartoon of the right-side up assay plate). Azide, a paralytic, is placed in each box. The placement of animals after they have run for 16-18 hours was noted. The Chemotaxis Index refers to how well a trained population retains memory to avoid butanone (refer to first formula), and the Learning Index reflects how much each strain has learned to avoid butanone after training (refer to second formula). **Figure 1b.Memory dynamics demonstrate the difference in memory consolidation between *osm-9(ky10)* and WT.**The Learning Index reflects how much each strain has learned to avoid butanone and how that memory has persisted over time from right after two-cycle training to 120 minutes later. The LI for wild type (black dots) immediately after training (0’), after 30 minutes on food (30’) or 120 minutes on food (120’) is compared to that of *osm-9(ky10)* (pink dots). Each data point represents a population of > 50 animals and each point is an independent day of experimentation. **Figure 1c.*WT C. elegans* and *osm-9(ky10)* mutants learn the same immediately after training, but retain different amounts of information 120 minutes later.**Left, the Learning Indexes of *WT C. elegans* and *osm-9(ky10)* mutants right after recovering from odor training and right, the Learning Indexes of *WT C. elegans* and *osm-9(ky10)* mutants after 120 minutes of recovery on food plates. The Learning Index reflects how much each strain has learned to avoid butanone and how that memory has persisted over time from right after two-cycle training to 120 minutes later. Each dot on the graph represents an independent assay with greater than 50 worms. Each graph has SEM bars and paired two-tailed t-tests were run. **Figure 1d.*WT C. elegans* lose memory after 30 minutes on food, which they gain back after 120 minutes. *osm-9(ky10)* mutants, however, lose memory but do not gain it back.**Left, the differences of Learning Indexes between 0 and 30 minutes on food and right, between 30 and 120 minutes on food. Comparing WT (N2) and *osm-9(ky10)* to themselves at different time points to analyze how their memory changes. Each dot on the graph represents an independent assay with greater than 50 worms. The graph shows SEM bars and paired 1 way ANOVA with the Dunn’s correction for multiple comparisons.

After the last training, each population of worms is split into three groups for a total of six, each of which is washed in buffer. One group from each population is immediately transferred to a chemotaxis assay plate in order to understand how buffer-trained (control) and butanone-trained (odor) worms learned from their training (Figure 1A 3 top row). To quantify how well each population of *C. elegans* learns, first we calculate each training groups’ Chemotaxis Index (CI) where the number of worms attracted to the ethanol buffer is subtracted from the butanone and the resulting value is divided by the total number of *C. elegans* on the chemotaxis plate (Figure 1A, zoom box). This number reflects the population’s attraction to butanone. After calculating the (CI) of both training groups we can use the difference between the control CI and odor CI to assess the Learning Index (LI) of the population. The LI is achieved where the CI value of the butanone trained group is subtracted from the buffer control group. The LI tells us how well the worms learned where a greater value indicates a larger difference between the two groups’ attraction to butanone.

In order to assess the changes in memory as the animals recover from starvation on food, we place animals on food and test their learning (LI) at intervals of 30 and 120 minutes (Figure 1A, 3 bottom two rows). Surprisingly, we found that though animals learn to avoid butanone immediately after training, they seem to lose this aversion following 30 minutes of recovery on food (Figure 1B, wildtype, black data points). However, by 120 minutes of recovery the aversion reappears. This reveals that memory consolidation in *C. elegans* is a discontinuous process that may evolve in two phases.

We asked if the *osm-9(ky10)* mutants differed from Wild Type *C. elegans* in the dynamics of their memory consolidation (Figure 1B, *osm-9(ky10),* pink data points). We found that the *osm-9(ky10)* mutants and Wild Type *C. elegans* LI were not significantly different immediately after the training (Figure 1B and 1C, left bars). By 120 minutes, the LIs of *osm-9(ky10)* mutants was significantly lower than Wild Type (Figure 1C, right bars), indicating that without the OSM-9 protein, animals are unable to regain memory.

To quantify the changes in memory at each stage in memory consolidation - the stage at which it is lost i.e. in the first 30 minutes and the stage during which it is regained (30 to 120 minutes) - we compared the difference in LIs at 0 and 30 minutes (Figure 1D, left bars) and those at 30 and 120 minutes (Figure 1D, right bars). We found that for the Wild Types, the change in LI was negative in the first 30 minutes and this was significantly different from the positive change in LI from 30 minutes to 120 (p-value of <0.01). To understand if the role for OSM-9 in this change from loss to gain of memory is significant, we compared each rate of change at each stage with the genotype using an ANOVA. We found that there was not a significant difference between both Wild Type *C. elegans* and *osm-9(ky10)* mutants for the difference of Learning Indexes between 0 and 30 minutes. This tells us that Wild Type *C. elegans* and *osm-9(ky10)* mutants lose the same amount of memory after 30 minutes and that the OSM-9 protein does not cause a difference in memory consolidation between immediate learning and the formation of short term memory.

The test revealed that there is no significant difference for the two time intervals 0 to 30 minutes and 30 to 120 minutes for *osm-9(ky10)* mutants (Figure 1D 2nd and 4th bars). This tells us that *osm-9(ky10)* mutants lose memory (the 0’ to 30’ LI change is negative, Figure 1D, 2nd bar) but do not regain it (the 30’ to 120’ LI change is not significantly different compared to the 0’ 30’ LI change, Figure 1D, 4th bar). Ultimately, this indicates that the loss of the OSM-9 protein affects the memory dynamics of *C. elegans* between the formation of short term memory and long term memory.

## Discussion

Spaced training – where starvation, the Unconditioned Stimulus (US), is paired with butanone, the Conditioned Stimulus (CS), and interleaved with feeding in the absence of the CS – induces memory that lasts for at least 24 hours. Previously, (Chandra, Farah, Munoz Lobato et al., 2023) we showed that this memory depends on sleep for its consolidation. In this study we show that this memory consolidation is dynamic in that it seems to disappear after 30 minutes and then recover after 120 minutes of recovery on food. Further, we show that OSM-9, the TRPV5/6 channel protein, is not required for initial learning but is required for memory after a recovery on food for 120 minutes. Upon close examination of the memory dynamics, OSM-9 is not required for the 30 minute food-triggered “forgetting” but is required for the recovery of memory between 30 and 120 minutes on food.

### OSM-9 is not impaired for learning after spaced training

We show here with two-cycle spaced training that *osm-9* mutants can learn the same as wild-types. Even though OSM-9 is expressed in sensory neurons AWA, ADL, ADF and ASH, (Benedetti et al., BioRxiv 2022) all of which respond to food and its removal (Ferkey et al., 2021), the fact that *osm-9* mutants learn makes it likely that they still perceive the US. Additionally, both *osm-9* mutants and wild types respond to food as a memory loss trigger. Thus, instead of being required for food sensation, OSM-9 may be involved with memory formation at a subsequent time. It could instead be required for neuromodulation or circuit dynamics that are triggered by these neurons in the process of memory consolidation.

### Memory dynamics - disappearing memory

Similar studies in other systems have revealed that long term memory can appear discontinuous (Kendall et al., 2014, Davis, 2023). In mice and human studies, memory becomes labile during the two hour period where sleep is required for consolidation immediately after being encoded. Likewise, in flies, transitions between short, intermediate and long term memory can appear abrupt. In mammals, the sleep-dependent transition reflects movement of the memory engram from the hippocampus to other regions of the brain.

The reasons for this discontinuity could reflect the transition between short and longer term memory which may require a rest period where signaling molecules replenish, such as the case of *drosophila* learning where ERK kinase levels are decreased by neuronal activity gradually rising during rest (Pangani et al., 2009). We do not have evidence for this yet in *C. elegans* but OSM-9 may be required for the recovery. In Benedetti et al., 2022, we show that OSM-9 mutants sleep just as well as wild types. Therefore, if OSM-9 is required for building up a memory factor, it is required regardless of whether the animal sleeps.

Another reason that memory may appear discontinuous we posit is that when memory is initiated, forgetting is also engaged. The disappearance of memory that we observed could result from the difference between the rate of the forgetting process relative to that of the long-term memory. Thus, the forgetting and long-term memory pathways may ramp up as short term memory declines but if the forgetting pathway ramps up more steeply, memory would seem to disappear until the increase in long term memory overtakes it. This forgetting pathway in fly olfactory learning (Davis 2023) and diacetyl learning in *C. elegans* (Hadziselimovic*elzmovic* et al., 2014) is mediated by actin reorganizing factors that prune synapses. In *osm-9* mutants, the forgetting may remain intact as they lose memory and may be over-active since they don’t seem to regain it back as they normally should. Or, *osm-9* mutants may be defective in the ramping up of long-term memory.

### Forgetting

OSM-9 loss may promote forgetting if OSM-9 is required to silence the “forgetting” cells or it may block cells that build a long-term memory. To establish how OSM-9 is acting, it will be important for us to understand which cells it acts in for long-term memory. To this end, we have identified four sensory neurons that are required to express OSM-9 in order for animals to regain memory (Bendetti et al., 2022). However, we still do not know how loss of OSM-9 affects these cells’ activity in response to either training or re-feeding after training.

### Neuromodulators

In *drosophila*, neuromodulators have been shown to be important in promoting memory as well as forgetting. Both OSM-9 expressing ADF and ADL neurons secrete neuromodulators. ADF secretes serotonin as well as the FMRFamide-like peptide FLP-6, the food triggered insulin-like peptide INS-1 and the neuropeptide-like peptide, NLP-3 while ADL secretes five neuropeptides (FLP-4, FLP-21, NLP-7, NLP-8 and NLP-10) (Li and Kim, 2008; Altun et al., 2009; Bhattacharya, Abhishek et al. 2019). Loss of OSM-9 could block or even enhance neuropeptide secretion. Thus, visual reporters for neuropeptides and serotonin could reveal whether loss of OSM-9 affects their secretion from these neurons. Loss of function mutants for each neuromodulator exist and these mutants could be tested for their memory dynamics to understand if they phenocopy the memory dynamics of *osm-9* mutants. Likewise, over-expression of these peptides or exogenous application of serotonin may alleviate or exacerbate the *osm-9* mutant’s memory deficits. Such studies would help establish the molecular mechanisms by which OSM-9 affects memory dynamics and open a window into how long-term memory is consolidated in the most anatomically simple nervous system that we know exhibits long-lasting sleep-dependent memory.

### Caveats to the study

This study is based on analysis of behavior and mutants. The difference between wild type and *osm-9* mutant learning was not significant but it trended towards osm-9 learning slightly less than the wild type. The observations are robust and rigorous but their interpretations are tentative as the next studies with more real time tools such as calcium imaging and cell specific and timing specific expression will need to be employed to test the hypotheses put forth.

## Methods

### Chenotaxis Assay Training

Using the chemotaxis assay from Bargmann et al., 1993, we performed two-cycle training assays on the *C. elegans*. 5 days before the assay, 4 Larval stage 4 (L4) worms are picked and transferred onto NGM plates that are seeded with OP50. These worms are to be undisturbed for 5 days at 20 degrees Celsius to ensure that the population of *C. elegans* are one-day old adults.

On the day of the assay, the plates will be observed for any bacterial or fungal contaminations; any plates with forms of contamination will not be used for the chemotaxis assay. *C. elegans* are then washed off the plates using S. Basal buffer into eppendorf tubes specific for their strain.

Worms are washed on the plate for a minimum of 5 washes until they are placed into the eppendorf tubes for an additional 3 washes. Once the worms are as evenly distributed between tubes, they will be conditioned to either a S. Basal buffer or 2-butatanone solution. The S. Basal buffer includes 0.1M NaCl and 0.05M K3PO4, maintaining a pH of 6.0. The 2-butanone solution is diluted with the S. Basal buffer with a ratio of 1 microliter to 1 milliliter. The worms are then spun on a rotisserie in their respective training odors – the buffer or butanone solution – for a total of 80 minutes. Afterwards, the worms are washed three times and then spun on a rotisserie for 30 minutes in an OP50 diluted solution. This cycle of going from training with buffers to being on a rotisserie with OP50 will be done for a total of 2 odor-treatment cycles and 1 OP50 cycle in between. Once the worms are washed for a total of 3 times with the S. Basal buffer, the worms will be split into three for two of the groups to be recovered at 30 minutes and 120 minutes. When the three final washes occur after the last training cycle, worms will be placed on their respective plates: the first group of worms will go onto chemotaxis assay plates while the other two groups are transferred onto recovery plates with food and chemotaxis assay plates later on.

### Chemotaxis Assay Plates for Observation

Chemotaxis plates are made by combining 200 milliliters of distilled water to 3.2 grams of Difco Bacto agar and boiling the mixture until all of the agar beads are melted. Once the agar mixture has cooled to 55 degrees Celcius, 1 milliliter of 1M K3PO4, 200 microliters of 1M CaCl2 and 200 microliters of 1M MgSO4 are pipetted to this mixture, ensuring that no precipitate has formed. Then, the mixture is pipetted at 9 milliliters per 10 centimeter Petri dish and left to stand for 15 minutes to solidify. Chemotaxis plates are to be expected to be made in the first training cycle to ensure that the plates are ready for use once the last remaining washes are done and during the recovery time periods.

Once the plates have solidified, square odor areas and an origin will be drawn and labeled: one square odor area labeled butanone and the other labeled ethanol. Then, each point source will contain 1 microliter of 1M Sodium azide and 1 microliter of each odor will go to each point source with account to their labeling. The purpose of the Sodium azide is to paralyze the worms at their respective odor areas for observation. Worms designated to be evaluated right after the training are then placed at the origin and liquid is removed using a Kim Wipe, allowing the *C. elegans* to freely roam around on the chemotaxis plate for at least 2 hours.Worms recovering at 30 and 120 minutes will then be observed and placed on plates when the time comes. Once all of the chemotaxis plates are observed, the chemotaxis index and the learning index will be calculated.

### Statistical Analysis

All data sets were subjected to the Shapiro-Wilk normality test before performing further analysis. For comparing 0’ minutes to 120’ minutes, if all of the datasets were normally distributed (Figure 1C), then parametric paired two-tailed t-tests were performed. If the datasets appear to be non-normally distributed, then the Krushka-Wallis test was performed. If p>0.05, then no further analysis was performed. Labels to compare data in the data set used depended on the p values calculated: p value <0.001 will be labeled with ***, p value <0.01 will be labeled with **, p-value <0.05 will be labeled with *, and NS will indicate that the p value >0.05.

For comparing the differences of LIs at two time points, if all of the datasets were normally distributed (Figure 1D), then parametric paired one-way ANOVA were performed, followed by Dunn’s multiple comparison test. If the datasets appear to be non-normally distributed, then a non-parametric unpaired one-way ANOVA was performed, followed by Dunn’s multiple comparison test.

All graphs show S.E.M. Graphpad Prism was used for all of the statistical tests. Each data point (the grey dots) on the graph indicate one population that was run on independent days.

**Table 1:**
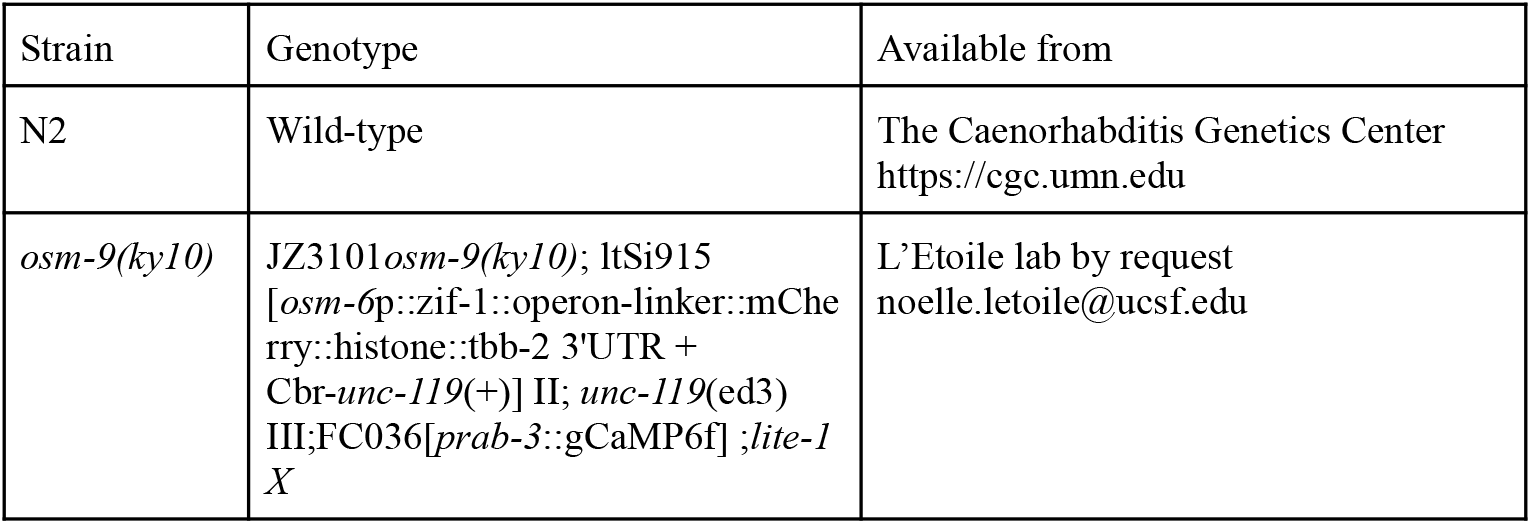
Strains Used.

## Acknowledgements

Trisha Yee for assistance during experiments.

Kevin Daigle for picking worms for assays and double checking contamination of assay plates. Dr. Laura Persson for insight into the conceptualization of the project.

## Funding

NIM Lowell High School Science Research Program Internship (Lowell Alumni Association) FMS F31A137849

NDL NINDS R01NS087544

## Author Contributions

Nikki Ma: Methodology, Investigation, Formal Analysis, Writing - original draft, Funding acquisition, Conceptualization

Fatema Mashel Saifuddin: Funding acquisition, Conceptualization Nando Munoz Lobato: Conceptualization

Noelle L’Etoile: Supervision, Methodology, Writing - original draft, review & editing, Funding acquisition, Conceptualization

